# ComBat-Predict enhances generalizability of neuroimaging models to new sites

**DOI:** 10.1101/2025.08.21.671401

**Authors:** Yao Xin, Margaret Gardner, Nick Tustison, Philip Cook, James Gee, Andreana Benitez, Jens H. Jensen, Alzheimer’s Disease Neuroimaging Initiative, Lifespan Brain Chart Consortium, Richard Bethlehem, Jakob Seidlitz, Aaron F. Alexander-Bloch, Andrew An Chen

## Abstract

Neuroimaging is vital in quantifying brain atrophy due to typical aging and due to neurodegenerative diseases. To collect large samples necessary to model lifespan brain development, research consortiums aggregate images acquired across multiple study sites. Previous studies have demonstrated that this multi-site study design can lead to site-related bias, necessitating harmonization of these “site effects”. However, current methodologies are unable to generalize to new sites outside the original harmonized sample, limiting translation to new sites or clinical practice. Here, we propose a method called ComBat-Predict (CB-Predict) building upon the ComBat method for site effect adjustment, which extends to data from a new site with smaller sample sizes and unknown site effects. In data from the Alzheimer’s Disease Neuroimaging Initiative (ADNI), our proposed method mitigates bias and yields high accuracy in predicting cortical thickness measures when generalizing the model to new data. Furthermore, we demonstrate that our proposed harmonization method can reduce site-related variance in centile scores estimated using data from the Lifespan Brain Chart Consortium (LBCC). Altogether, our results demonstrate that CB-Predict effectively harmonizes new sites and thereby enables effective translation of neuroimaging models to additional samples.

## Introduction

Combining data from studies conducted at different locations can be used to increase sample size thereby enhancing statistical power. This need arises when individual studies are limited by practical or logistical constraints on sample size, or when researchers aim to aggregate data collected across various environments to improve generalizability (Solanes et al., 2021). However, pooling data sets is generally considered inappropriate without prior adjustment for differences between batches or sites, which can introduce systematic measurement errors (Chen, Luo, et al., 2022; Fortin et al., 2016, Fortin et al., 2017, Fortin et al., 2018; Kruggel et al., 2010; Solanes et al., 2021). These non-biological variations, referred to as site or scanner effects depending on the context, can arise from variability in scanner hardware, acquisition parameters, and even processing pipelines (Reig et al., 2009; Kruggel et al., 2010; Chen, Beer, et al., 2022). Studies have found that site effects are present in various modalities including structural magnetic resonance imaging (MRI), diffusion tensor imaging and functional MRI (Fortin et al., 2017; Yu et al., 2018). Critically, failure to address these site effects can significantly decrease statistical power (Fortin et al., 2016, Fortin et al., 2017), reduce the reliability of measurement, mask biologically meaningful associations, and potentially introduce bias or spurious findings (Yamashita et al., 2019; Chen, Beer, et al., 2022).

In parallel, normative modeling in neuroimaging has emerged as a powerful approach for evaluating individual brain characteristics against population reference standards, allowing researchers and clinicians to quantify how an individual’s brain metrics deviate from expected patterns in the population (Gage et al., 2024). While the need for such standardized reference tools, akin to growth charts, has long been recognized (Cole, 2012), early efforts often utilized smaller datasets or specific age ranges (Ber et al., 2017; Bozek et al., 2023; Dong et al., 2020). A landmark achievement involved aggregation of over 100,000 scans to create robust lifespan brain charts, which are used to estimate centile scores relative to age- and sex-matched peers (Bethlehem et al., 2022). This approach has rapidly gained prominence, fueled by the development of increasingly large reference cohorts (Córdova-Palomera et al., 2021; Nobis et al., 2019; Rutherford et al., 2023), dedicated repositories, established statistical modeling frameworks (e.g., generalized additive models for locations scale and shape (GAMLSS), recommended by the World Health Organization), and accessible tools facilitating their application (Rutherford et al., 2023). Furthermore, studies have demonstrated that features derived from normative modeling can offer statistically significant advantages over raw data in clinical benchmarking tasks like group difference testing and classification (Rutherford et al., 2023). However, constructing normative models for the human lifespan requires massive sample sizes, only achievable through aggregation of MRI images across sites and studies. While GAMLSS-based normative models account for site effects, Bethlehem et al. noted that the unbiased and stable estimation of centile scores in new sites required a sample size above 100 (Bethlehem et al., 2022).

With the rise of international consortium neuroimaging studies and lifespan normative modeling, it has become increasingly important to distinguish site-related effects from genuine biological differences among the individuals studied (Bayer et al., 2022; Chen, Beer, et al., 2022; Gardner et al., 2024; Pomponio et al., 2020). Consequently, various statistical and machine learning methods have been proposed to address these site effects (Chen, Beer, et al., 2022; Fortin et al., 2016, Fortin et al., 2017, Fortin et al., 2018; Gardner et al., 2024; Hu et al., 2023; Pomponio et al., 2020). Some proposed deep learning approaches for mitigating site effects include CycleGAN (J.-Y. Zhu et al., 2017) and its variants, variational autoencoders (VAEs) including conditional VAEs and goal-specific VAEs (An et al., 2022; Moyer et al., 2020), and batch unlearning classifiers (Dinsdale et al., 2021; Hong et al., 2023). In addition, there are also methods based on latent factor modeling (Avalos-Pacheco et al., 2022; Fortin et al., 2016; Zhang et al., 2023) and on regression modeling using empirical Bayes (Johnson et al., 2007). Among methods using empirical Bayes shrinkage estimators, ComBat remains a widely-employed harmonization procedure owning to its strong performance in multiple MRI modalities, ease of interpretation, and robustness to small sample sizes (Fortin et al., 2016, Fortin et al., 2017, Fortin et al., 2018; Hu et al., 2023; Johnson et al., 2007).

Several limitations of ComBat have been addressed through recent methodological extensions. First, ComBat inadequately captures non-linear trajectories of brain development over the lifespan, prompting their development of ComBat-GAM, which integrates generalized additive models (Pomponio et al., 2020). Second, ComBat does not model covariate effects on variance, leading to biases in normative models. ComBatLS is a recent approach designed to preserve covariate-dependent variance structures alongside mean adjustments, yielded greater accuracy in centile scores from normative (Gardner et al., 2024). For machine learning studies, CovBat addresses multivariate site effects emphasizing that harmonizing correlations among neuroimaging measures is critical for downstream machine learning applications (Chen, Beer, et al., 2022). Additionally, Beer et al. (Beer et al., 2020) introduced Longitudinal ComBat, specifically tailored to harmonize repeated measures collected across multiple time points, addressing inherent challenges in longitudinal neuroimaging designs.

Nevertheless, key limitations of ComBat and its extensions are that data must be gathered at one site and harmonization must be re-applied each time a new cohort is added to the study. To address this challenge in clinical data sharing, Chen et al. introduced distributed ComBat (d-ComBat), enabling harmonization across distributed datasets while preserving data privacy (Chen, Luo, et al., 2022). However, d-ComBat requires multiple communications between study sites, leading to practical issues in implementation particularly with a large number of sites or clinical sites with strict communication restrictions. Recently, another group proposed an extension of ComBat-GAM to harmonize new sites to a reference site; however, their method still requires sharing of the reference site data which can be infeasible in normative modeling settings (A. H. Zhu et al., 2025).

The present study proposes an extension to the ComBat framework, ComBat-Predict (CB-Predict), allowing for the harmonization of new sites with unknown site effects. CB-Predict effectively harmonizes out-of-sample data, approximating standard ComBat results even with very small sample sizes (n < 5) or varying diagnostic compositions. In the Alzheimer’s Disease Neuroimaging Initiative (ADNI), which is previously shown to have notable site effects (Chen, Luo, et al., 2022), we demonstrate that applying CB-Predict improves the generalizability of predictive models to new sites. Further extending our analyses to data from the Lifespan Brain Chart Consortium (LBCC), our experiments show that CB-Predict can greatly improve generalizability of fitted normative models to the ADNI data, highlighting differences in centile scores between diagnostic groups.

## Methods

### ComBat framework

The location-and-scale-based adjustment method for site effects by Johnson et al. 2007^1^ has been shown to be effective in genomic and neuroimaging data, as a popular harmonization method robust to small sample sizes^2^ (Fortin et al., 2017, Fortin et al., 2018). To harmonize the data *Y*_*iJg*_ for site *i*, subject *j*, region *g*, the ComBat model is specified as

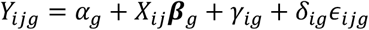

where *α*_*g*_ is the overall mean, *X*_*ij*_ is the design matrix, *β*_*g*_ is the vector of corresponding regression coefficients, *γ*_*ig*_ is the additive site effect, and *δ*_*ig*_ is the multiplicative site effect. The errors are assumed to be normally distributed, 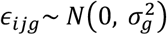 For identifiability, the additive site effects are constrained such that 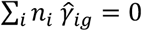. The Empirical Bayes method for estimating the additive site effect γ_ig_ and the multiplicative site effect δ_ig_ follows these steps:

Step 1: Standardize the data with least squares estimates for *α*_*g*_*, β*_*g*_,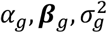:

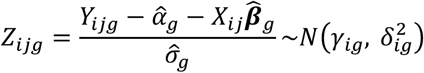

Step 2: Based on conjugate priors: 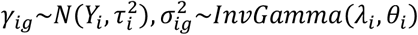 and method of moments estimates 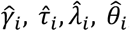 obtain the posterior estimates 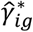 and 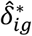

Step 3: Adjust the data:

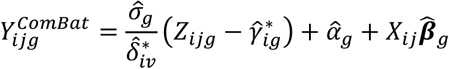

### ComBat-Predict extension

To extend the ComBat framework to harmonize new sites, we propose ComBat-Predict (CB-predict) which can estimate adjustment parameters in the new site without the need to pool it with data from the original sites.

Suppose we have *K* original sites and we fit ComBat on those sites *i* = 1, 2, … , *K*. Our goal is to estimate harmonization parameters for a new site *K* + 1. In Step 2 of the ComBat algorithm, the method of moments estimates of the parameters for *γ*_*i*_ , *τ*_*i*_ , *λ*_*i*_ and *θ*_*i*_ in site *i* are obtained via

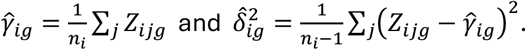

When extending to the new site *K* + 1, we first standardize the new site data using the original ComBat model estimates 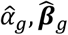 and 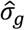 to obtain *Z(*_*K*+1*)jg*_. Then, method of moments estimates can be derived via

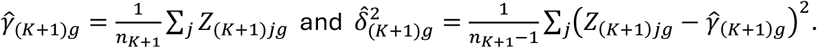

After subsequent Empirical Bayes estimation, we obtain posterior estimates of the site effects 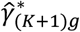 and 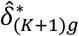 Finally, we harmonize the data in site *K* + 1.

In a more general setting, the standardized of *Z(*_*K*+1*)jg*_ can also be derived using a broader approach than described above. For example, in the ComBat extension incorporating Generalized Additive Models (GAM), the standardized value is computed as:

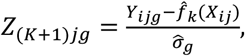

where 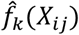 represents a GAM-based covariate function estimate, allowing non-linear modeling of effects such as age, sex, and other variables (Pomponio et al., 2020).

Furthermore, CB-Predict can extend ComBatLS through replacement of 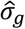 with a subject-specific standard deviation estimate 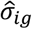 described previously (Gardner et al., 2024).

Our method is implemented as part of the ComBatFamily R package as *predict*.*comfam* and publicly accessible via Github: https://github.com/andy1764/ComBatFamily.

### ADNI data processing

The ADNI dataset in these analyses were originally processed for Tustison et al., 2019 and also previously described in Chen et al., 2021 and Beer et al., 2020. From 833 ADNI1 (http://adni.loni.usc.edu/) participants, cortical thicknesses for 62 brain regions defined by the Desikan-Killinary-Tourville atlas (Klein & Tourville, 2012) were obtained in the downstream of MRI scans processed by using the ANTs longitudinal single-subject template pipeline, with code available on GitHub (https://github.com/ntustison/CrossLong/). In summary, T1-weighted images from the ADNI-1 database were acquired using MPRAGE sequences across different scanners. For each subject, a within-subject template from all image time points was created, and all timepoint images were rigidly aligned to it before undergoing a standardized cortical thickness pipeline involving brain extraction, denoising, bias correction, tissue segmentation, and thickness estimation. Baseline cortical thickness values (62 regions) were extracted, and scanner information was determined from Digital Imaging and Communications in Medicine (DICOM) headers (Beer et al., 2020; Chen, Luo, et al., 2022).

Categorizing the scanners by each unique combination of study site, manufacturer, scanner model, coil type, and magnetic strength, we define 104 “sites” in this dataset for the current study. After removing sites with sample sizes smaller than 3 to ensure feasibility of harmonization, 505 subjects from 64 sites remain, with the sample size within each site ranging from 3 to 20. The majority of scans were acquired at 1.5T field strength, with 32 subjects from 8 sites scanned at 3T.

### LBCC data preprocessing and processing

The Lifespan Brain Chart Consortium (LBCC) comprises a large-scale collection of 1,239,894 structural MRI scans from 101,457 human participants, aggregated from over 100 globally diverse sources and spanning the entire human lifespan. Detailed descriptions of the contributing primary studies, dataset composition, and diagnostic classification criteria are provided by Bethlehem *et al*. (2022). Gardner *et al*. (2024) utilized a subset of the LBCC compiled from 60 of these studies, removed scans conducted on individuals below 3 years old, and randomly selected one single scan for subjects with multiple scans. Further quality control was performed by excluding scans that failed to meet an adaptive threshold based on the Euler index. The resulting sample included 55,237 high-quality scans obtained from 202 imaging sites across 51 studies.

Using this processed data, we create a subset focused on healthy adults drawn from 49 studies. Subsequently, we exclude participants based on specific criteria: 188 individuals are removed due to any missing cortical thickness measurements, and 3 individuals are removed as they belong to 2 studies with fewer than 3 participants. Additionally, the entire ADNI study is excluded to prevent overlap with the ADNI dataset. These steps yield a final sample of 12,396 subjects originating from 45 different studies. For analyses considering site effects, the primary study is used as the ‘site’ identifier. The sample sizes for these sites (studies) vary, ranging from 5 to 1,132 participants. The original studies employed scanners with varying magnetic field strengths (1.5T, 3T, and others), with detailed specifications provided in the original publication.”(Bethlehem et al., 2022)

Both ADNI and each primary study who’s sample is included in the LBCC obtained written informed consent from all participants.

### Evaluation of CB-predict performance in a leave-one-site-out experiment

To assess performance of CB-Predict for harmonization of new sites, we leave each of the 64 ADNI sites out and fit a ComBat model on the remaining 63 sites. The cortical thickness measures in the left-out site are then harmonized with CB-Predict. To quantify the difference between CB-Predict and ComBat, we compute root mean squared error (RMSE) in the left-out site comparing harmonization using CB-Predict and ComBat including all 64 sites. D-ComBat is equivalent to ComBat including all sites (Chen, Luo, et al., 2022); therefore, we are effectively comparing CB-Predict versus ComBat and d-ComBat in our experiment.

### Generalizability of predictive models to a new site

To emulate the real-world scenario where a pre-established predictive model is fit on private data, we again use a leave-one-site-out experimental design and fit a multivariate linear model on the ComBat-harmonized cortical thickness measures in 63 of the 64 sites. The linear model regressed each cortical thickness on age, sex, and diagnosis (categorized as Alzheimer’s disease, late mild cognitive impairment, or cognitively normal). In the left-out site, we then predict the 62 regions of cortical thickness measures from age, sex and diagnosis. Here, our goal is to compare the predicted cortical thickness values to the unharmonized observed values and the CB-Predict-harmonized values.

Harmonizing the testing data independently with CB-Predict (as described in **Results section 3**), we obtain RMSE between ComBat-Predict harmonized data and the linear model prediction. Then we compare the RMSEs to the those between the unharmonized measures and the predicted values.

### Generalizability of normative models to a new site

To simulate applying a pre-trained normative model while accounting for novel site effects, we first established a reference model using 62 cortical thickness measures from the LBCC dataset. These LBCC measures were harmonized using ComBat (Johnson et al., 2007), treating study as the site variable and including age and sex as biological covariates. Subsequently, separate linear models were fitted for each of the 62 harmonized cortical thickness measures, modeling the relationship with age and sex. Analyses conducted are under the assumption of normally distributed residuals and a linear relationship between each outcome with age. These models provided age-and-sex-specific predicted means and residual standard deviations, forming the basis of our normative reference. Assuming a normal distribution with this normative reference, a cumulated distribution percentile was derived as the normative centile score for each individual in each region.

Next, we apply this LBCC-based GAMLSS model to the independent ADNI dataset. We only include ADNI sites containing at least 3 healthy control subjects to ensure feasibility of harmonization fitting. Our final subsample consists of 300 individuals from 30 ADNI sites. We apply our fitted GAMLSS model to obtain centile scores from the unharmonized cortical thickness data. Then, we apply CB-Predict to the 30 ADNI sites, using the harmonization fit from the LBCC data. Using the ComBat model parameters originally estimated from the LBCC reference data, we harmonized the ADNI cortical thickness measures via the CB-Predict framework, leveraging the control subjects within the target ADNI sites to estimate the necessary site-specific adjustments. For each ADNI subject, their age-and-sex-specific expected mean and standard deviation were then predicted using the corresponding linear models trained on the harmonized LBCC data. Normative centile scores were calculated for each testing set individual by comparing their *harmonized* cortical thickness value to their predicted distribution (mean and SD) using a normal cumulative distribution function. To specifically isolate the impact of applying harmonization to the test data, we performed a parallel calculation generating centile scores using the subjects’ *original, unharmonized* cortical thickness values against the same normative reference from the harmonized LBCC training data.

Finally, to assess if harmonization reduced the heterogeneity attributable to study site between the healthy reference (LBCC) and test (ADNI) cohorts, we compared their centile score distributions. For each of the 62 regions, we utilize Wilcoxon rank-sum tests to test for differences between healthy LBCC controls and healthy ADNI controls, using both harmonized and unharmonized ADNI scores separately. Rank biserial correlation quantified the effect size of these distributional differences, allowing comparison of heterogeneity reduction achieved by harmonization across regions.

## Results

### ComBat-Predict Consistently Harmonizes Data from Novel Sites

To evaluate the performance of ComBat-Predict (CB-Predict) in harmonizing data from previously unseen sites, we conducted a leave-one-site-out cross-validation using the 64 ADNI sites. The accuracy of CB-Predict was assessed by comparing its harmonization output for the held-out site against the harmonization achieved by a standard ComBat model applied to all 64 sites simultaneously, using the Root Mean Squared Error (RMSE) averaged across subjects within the left-out site as the primary metric (Figure 1). Across the 63 evaluated sites (site 64 served as the reference batch and was excluded), the discrepancies between the CB-Predict harmonization and the ComBat harmonization were consistently low. Figure 1a demonstrates that the RMSE remains low even when the left-out site contains a small number of subjects, with sample sizes ranging from 3 to 20. Additionally, Figure 1b indicates that this effective harmonization by CB-Predict was robust to variations in the proportion of individuals with an Alzheimer’s disease diagnosis within the left-out site. These findings suggest that CB-Predict effectively approximates the harmonization achieved by standard ComBat when applied to novel sites, regardless of common variations in site sample size or diagnostic composition.

**Figure 1.**
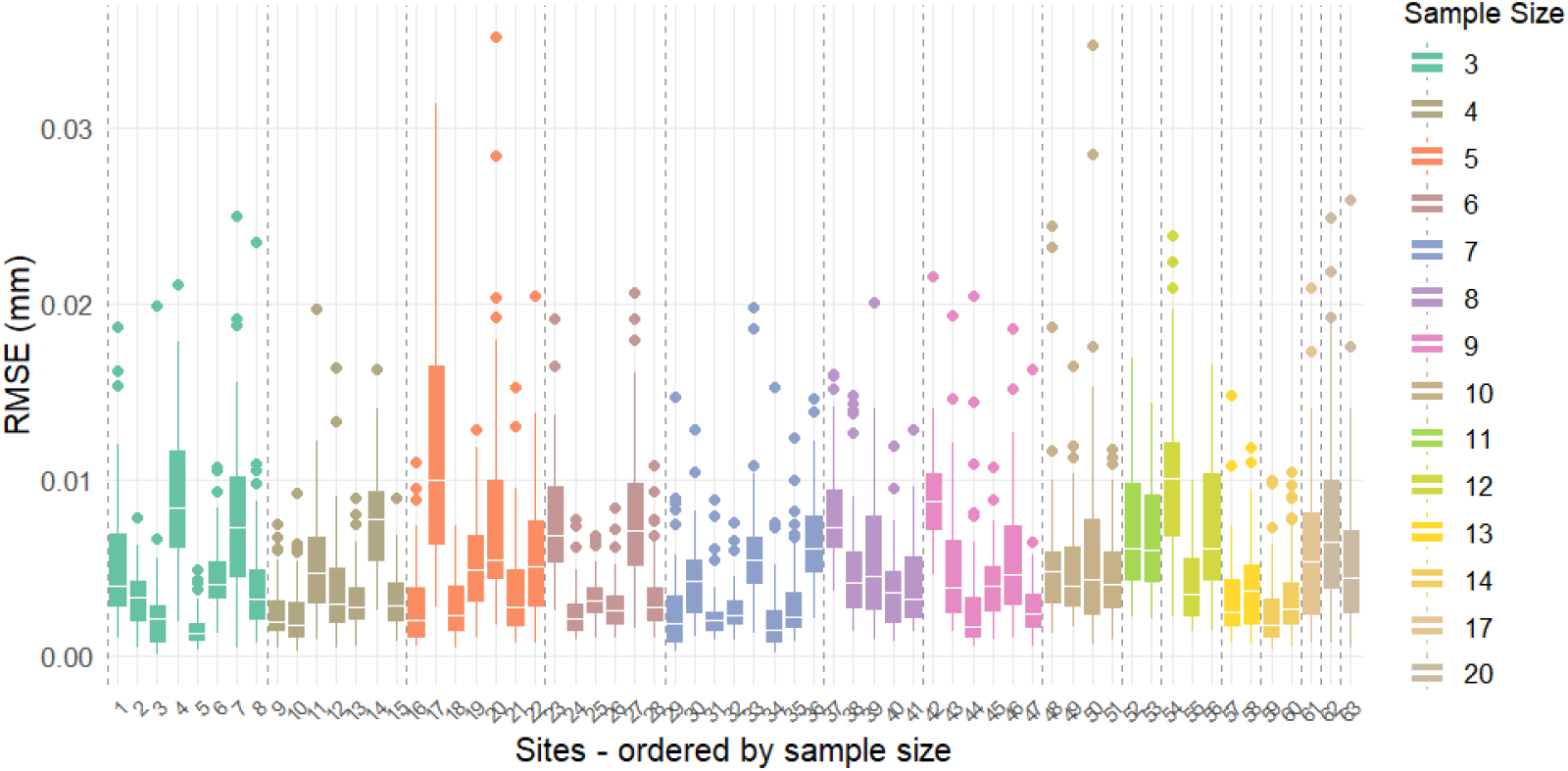
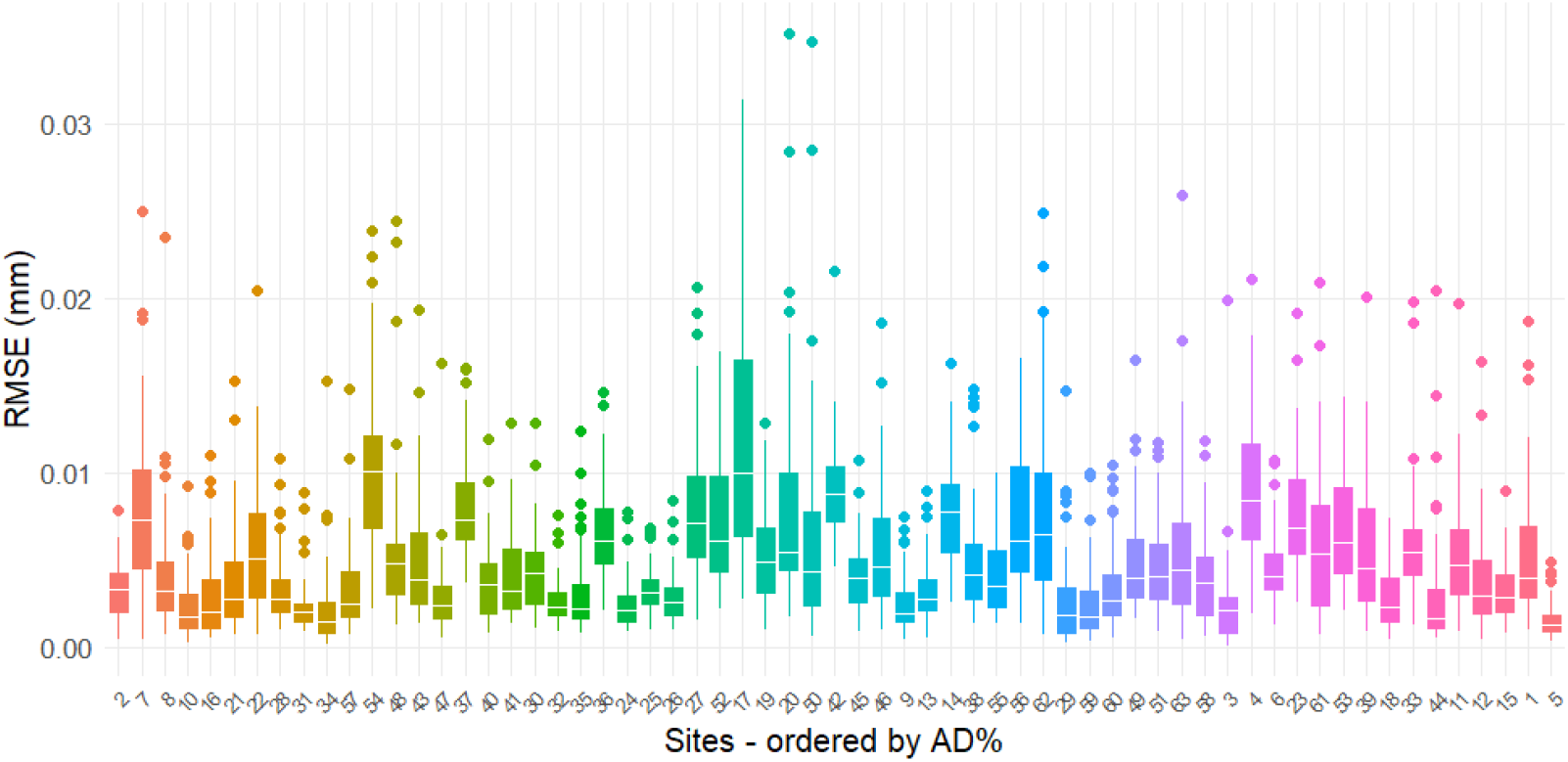
ComBat-Predict effectively harmonizes new sites, evaluated by leaving one site out. The difference between the in and out-of-sample harmonization were low in for all 1-63 sites (site 64 is used as a reference site and thus excluded). **a)** The mean squared errors (MSEs) are consistently low across different sample sizes in the testing sites, ranging from 3 to 20. **b)** The MSEs were consistently low across different proportions of Alzheimer’s disease diagnosis in the testing sites.

### CB-Predict Improves Predictive Model Generalization to New Sites

We next investigated whether applying CB-Predict harmonization to data from a novel site could improve the generalization performance of a predictive model trained on previously harmonized data. Using the same leave-one-site-out framework, a multivariate linear model predicting 62 cortical thickness regions based on age, sex, and diagnosis was trained on ComBat-harmonized data from 63 ADNI sites. This model was then used to generate predictions for subjects on the held-out site. As illustrated in Figure 2, the Root Mean Squared Errors (RMSEs) between these predicted cortical thickness values and the observed values are calculated for each left-out site under two conditions: comparing predictions against the original, unharmonized observed data (grey boxplots) and against the observed data after harmonization with CB-Predict (green boxplots). Across the evaluated sites, the RMSEs are consistently lower for the harmonized data compared to the unharmonized data. This demonstrates that applying CB-Predict to harmonize data from a new site significantly enhances the accuracy of predictions derived from a model trained on harmonized data from different sources, thereby improving model generalizability.

**Figure 2.**
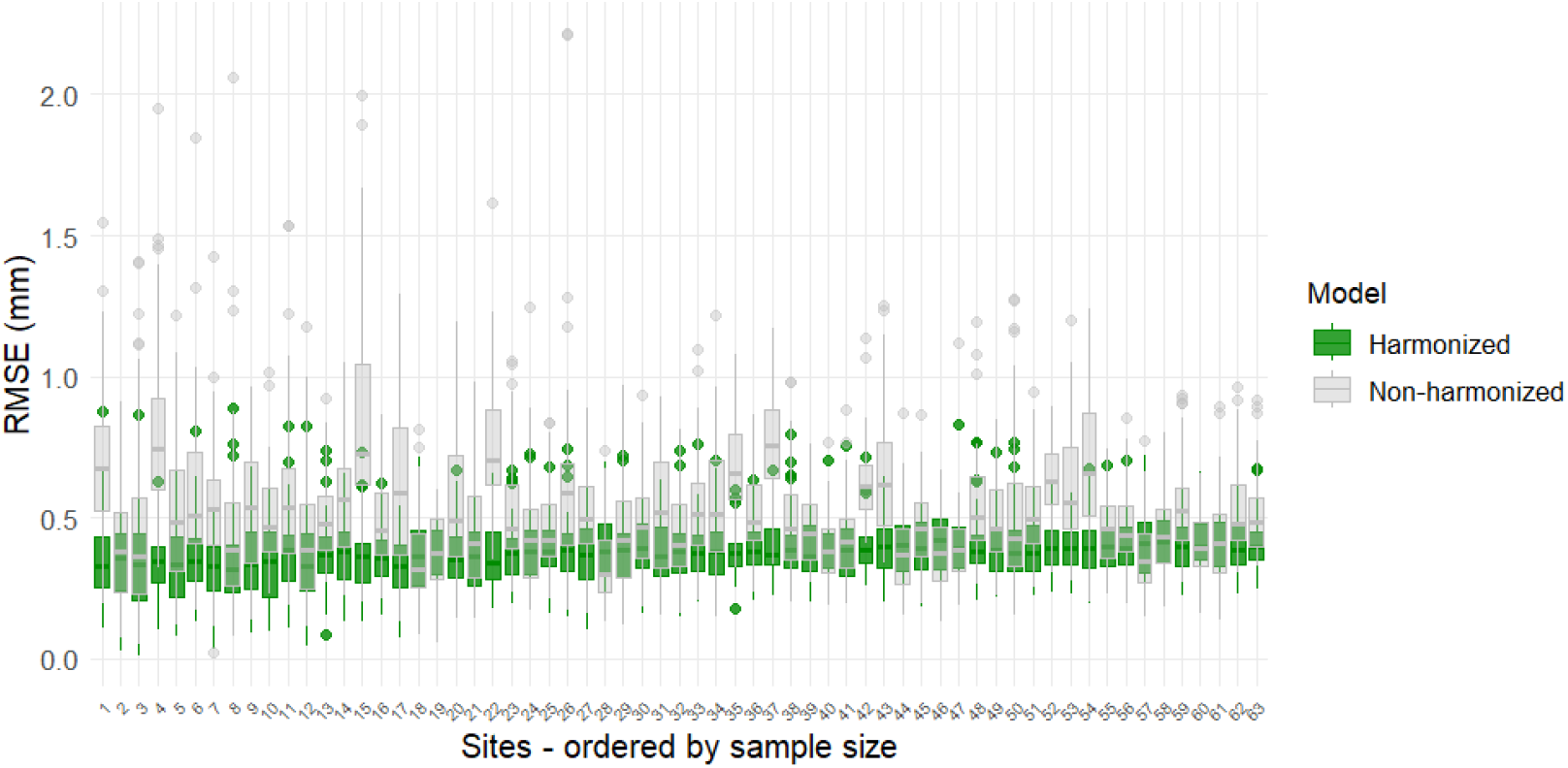
ComBat-Predict enables generalization of a fitted multivariate linear model to new sites. For the same left-out data in site *i*, compared to the MSEs between observed and predicted cortical thickness measures by the linear model based on age, sex and diagnosis (grey), the MSEs between ComBat-Predict harmonized and the same predicted outcomes (green) are notably lower.

### Harmonization Aligns Normative Scores Between Reference and Test Cohorts

To evaluate the application of the LBCC-derived normative model to the independent ADNI cohort and the impact of test-set harmonization, we generated centile scores for all 62 cortical thickness regions across ADNI diagnostic groups (CN, LMCI, AD), comparing them to the reference LBCC controls (Figure 3). When applying the normative model directly to the unharmonized ADNI data (Figure 3a), the resulting centile scores for the ADNI cognitively normal (CN) group (blue panels) showed marked deviations from the expected uniform distribution centered around 0.5 seen in the LBCC controls (grey panels). Across many regions, the ADNI CN scores exhibited systematic shifts and increased variance, indicative of residual site effects confounding the normative comparison. Applying the same normative model after harmonizing the ADNI data using CB-Predict (Figure 3b) resulted in a visibly improved alignment between the healthy groups. The centile scores for the harmonized ADNI CN group were more centered around 0.5 and displayed reduced region-specific biases, appearing more consistent with the reference LBCC control distribution. While the LMCI (cyan) and AD (orange) groups still showed deviations from the controls in both plots, reflecting pathological changes, the baseline comparison against the CN group in Figure 3b is less confounded by site effects compared to Figure 3a. This visual evidence, supported by quantitative comparisons (see next section), indicates that harmonizing the test data with CB-Predict substantially reduces site-related heterogeneity, leading to more comparable normative scores between the reference and test control groups.

**Figure 3.**
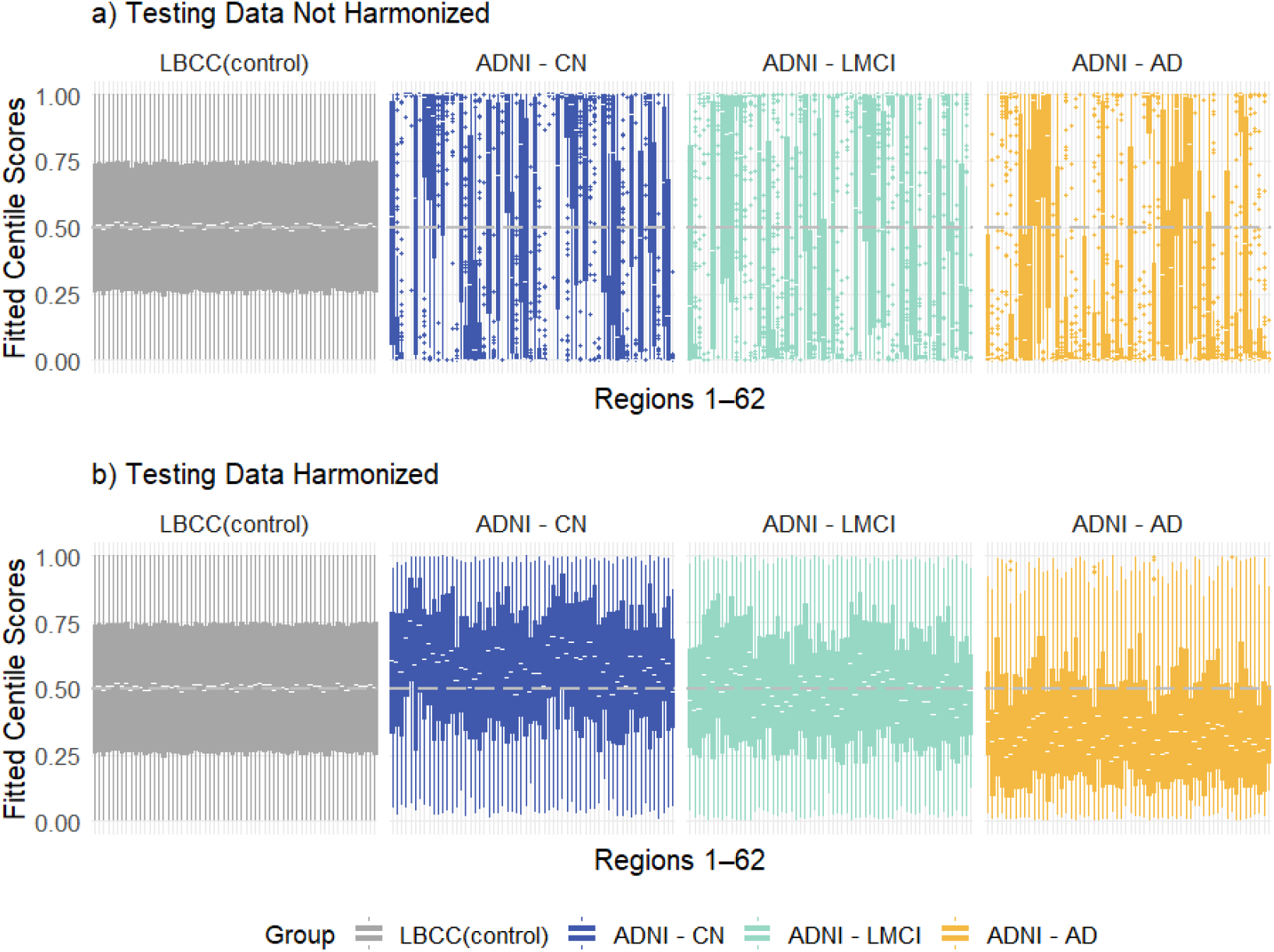
Comparison of cortical thickness centile scores across 62 regions from the reference LBCC controls and the ADNI test cohort (stratified by diagnosis: CN, LMCI, AD). (a) Centile scores derived from unharmonized ADNI cortical thickness data applied to the LBCC-trained normative model. (b) Centile scores derived after harmonizing the ADNI data using CB-Predict based on the LBCC ComBat fit, followed by application of the LBCC-trained normative model.

### 4. CB-Predict Harmonization Reduces Heterogeneity Between Healthy Control Groups

To quantitatively assess the reduction in heterogeneity between the healthy reference (LBCC) and test (ADNI) cohorts achieved by harmonization, we compared their centile score distributions using Wilcoxon rank-sum tests for each of the 62 cortical regions. The effect size of the difference between healthy LBCC controls and healthy ADNI controls was quantified using the rank biserial correlation (RBC), calculated separately using either the unharmonized or the CB-Predict harmonized ADNI centile scores (Figure 4). Figure 4 displays these RBC values for all regions. The RBC values derived from the unharmonized ADNI data (yellow points) show substantially larger magnitudes than those derived from the CB-Predict harmonized ADNI data (blue points). This demonstrates that applying CB-Predict harmonization significantly reduced the heterogeneity between the normative centile score distributions derived from healthy subjects in the novel study site and those from the larger reference population.

**Figure 4.**
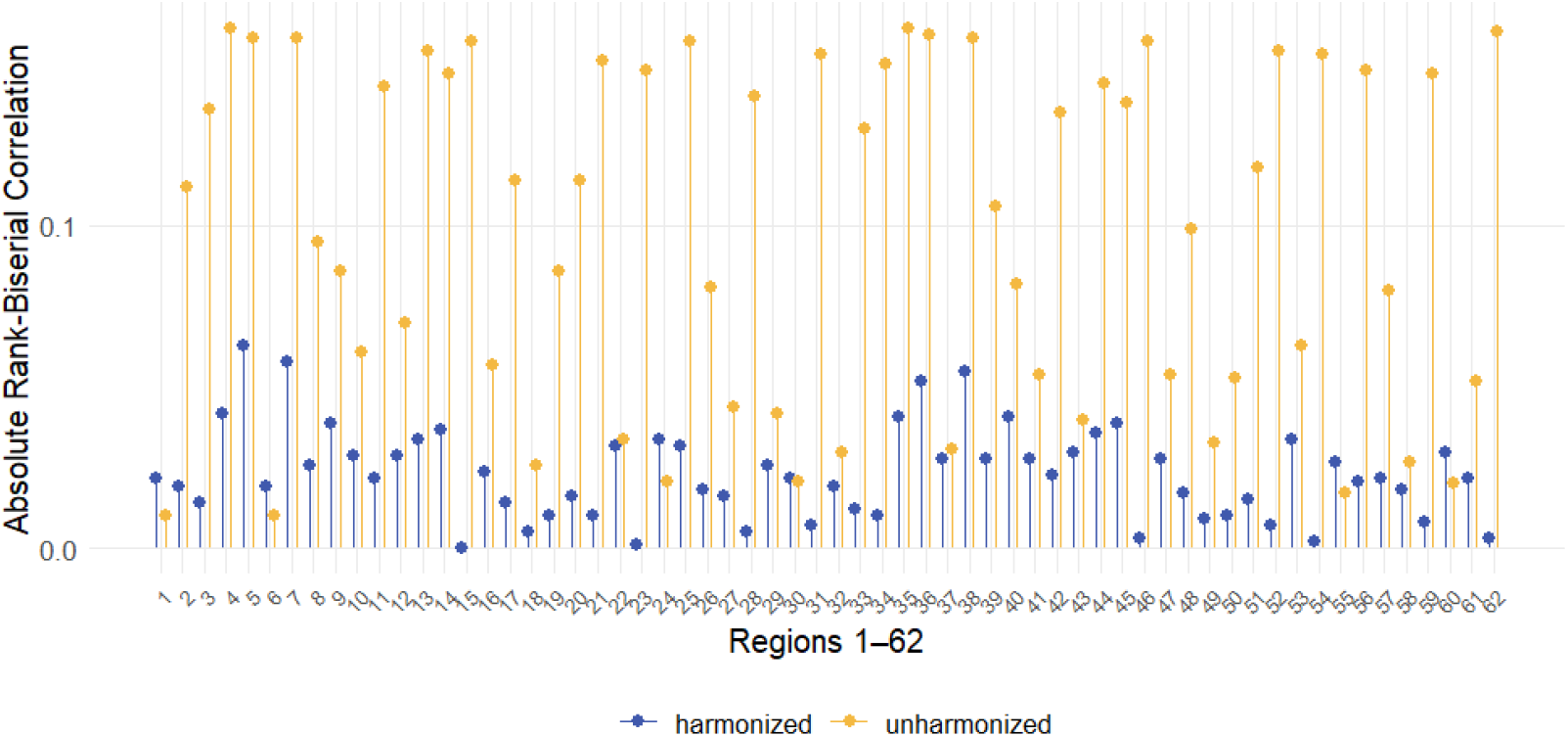
Harmonization reduces heterogeneity between healthy reference and test cohorts. Rank biserial correlation (RBC) effect sizes are shown for the comparison of centile score distributions between healthy LBCC controls and healthy ADNI controls across 62 cortical regions. Blue points represent effect sizes calculated using CB-Predict harmonized ADNI data, while yellow points represent effect sizes calculated using unharmonized ADNI data. Values closer to zero indicate greater similarity between the distributions of the two healthy control groups.

## Discussion

In the current study, we propose and validate ComBat-Predict (CB-Predict) for harmonization of new sites, without requiring access to data from the original sites. We evaluate the performance of CB-Predict by comparing to the conventional pooled sample ComBat harmonization and d-ComBat. Our leave-one-site-out experiment using ADNI data shows that CB-Predict consistently harmonizes data from previously unseen sites, effectively approximating the harmonization achieved by standard ComBat (Johnson et al., 2007) and d-ComBat (Chen, Luo, et al., 2022). We demonstrate this high performance is maintained across sites of varying sample size and subject diagnoses.

Furthermore, we validate CB-Predict for generalizability of predictive and normative models to a new site. In predictive modeling, our results indicate that applying CB-Predict to a new site lowers RMSE of predictive models trained on previously harmonized cortical thickness data. This experiment shows that applying CB-Predict enhances the accuracy and thus generalizability of pre-established model predictions to novel site data. In normative models, our experiments suggest that CB-Predict greatly improves comparability of new site centile scores with centile scores in the larger reference population. Examining a normative model trained on harmonized healthy data (LBCC) and applied to an independent cohort (ADNI), we observe reduced site-related heterogeneity and better alignment of centile scores in healthy controls from the LBCC and ADNI. Additionally, the distinctions in the centile scores of individuals with and without cognitive impairment become more observable.

However, we acknowledge several limitations that could be considered for future research. Firstly, the primary analyses were exclusively conducted on the ADNI1, which may not fully encompass the diverse demographic characteristics of populations outside of the United States. Future work could explore the method’s generalizability by evaluating its performance on other multi-site datasets including the international ENIGMA study (Thompson et al., 2020). Secondly, while the current method can be applied to other ComBat family extensions, such as ComBat-GAM and ComBatLS, its performance with these specific variants has not yet been evaluated in this study. Lastly, the study uses a single regression-based approach when generating the centile scores. Future research could explore the impact of CB-Predict harmonization on the robustness and interpretability of alternative normalization approaches including quantile regression (Koenker & Bassett, 1978; A. H. Zhu et al., 2025), hierarchical modeling (De Boer et al., 2024), and variational autoencoders (An et al., 2022; Moyer et al., 2020).

To conclude, we propose CB-Predict as an extension to the ComBat method, allowing for the harmonization of new sites with unknown site effects and small sample sizes. This method benefits imaging research practice by potentially avoiding the need to pool new site data with existing data for harmonization. With the current need for multisite studies in neuroimaging, CB-Predict’s ability to reduce site-related variance will contribute to robust analyses, to improvement in the generalizability of predictive models to new datasets, and to the alignment of normative scores between novel and reference cohorts.

## Data Availability

Data were obtained from the publicly available ADNI database (adni.loni.usc.edu) and the LBCC consortium dataset(brainchart.shinyapps.io/brainchart), both accessed under their respective data use agreements. Links to open datasets of LBCC are listed on https://github.com/brainchart/Lifespan. The investigators within the ADNI contributed to the design and implementation of the ADNI and/or provided data but did not participate in analysis or writing of this report. A complete listing of ADNI investigators can be found at https://adni.loni.usc.edu/wp-content/uploads/how_to_apply/ADNI_Acknowledgement_List.pdf.

## Code Availability

ComBat-Predict is implemented as part of the *ComBatFamily* R package as *predict*.*comfam* and publicly accessible via Github: https://github.com/andy1764/ComBatFamily.

## Author Contributions

**Conceptualization:** A.A.C. **Methodology:** A.A.C, Y.X. **Data curation:** M.G., J.G., R.B., J.S., A.A.B., N.T., P.C. **Statistical analysis:** Y.X. **Writing–original draft:** Y.X., A.A.C. **Writing– review and editing:** M.G., J.G., R.B., J.S., A.A.B., N.T., P.C., A.B., J.H.J. **Investigation:** ADNI, LBCC. **Supervision:** A.A.C. **Project administration:** A.A.C.

## Funding

A.A.Chen’s work was partially supported by National Institutes of Health (NIH) Grant R01MH123550.

The processing of the LBCC dataset was partially supported by NIH grants R01MH132934, R01MH133843, and R01MH134896. A complete list of funding sources that supported the collection and processing of data included in the LBCC can be found in Bethlehem et al, 2022.

## Declaration of competing interests

The authors have no conflicts of interest to declare.

## Acknowledgement

A complete list of acknowledgements associated with the LBCC dataset is available in Bethlehem et al, 2022.

## Disclosure

AAB, JMS, and RAB hold equity in Centile Bioscience. JMS and RAB are directors of Centile Bioscience.

## Notes

https://github.com/andy1764/ComBatFamily

### Competing Interest Statement

The authors have declared no competing interest.

## References

An, L., Chen, J., Chen, P., Zhang, C., He, T., Chen, C., Zhou, J. H., & Yeo, B. T. T. (2022). Goal-specific brain MRI harmonization. NeuroImage, 263, 119570. 10.1016/j.neuroimage.2022.119570

Avalos-Pacheco, A., Rossell, D., & Savage, R. S. (2022). Heterogeneous Large Datasets Integration Using Bayesian Factor Regression. Bayesian Analysis, 17(1), 33–66. 10.1214/20-BA1240

Bayer, J. M. M., Thompson, P. M., Ching, C. R. K., Liu, M., Chen, A., Panzenhagen, A. C., Jahanshad, N., Marquand, A., Schmaal, L., & Sämann, P. G. (2022). Site effects how-to and when: An overview of retrospective techniques to accommodate site effects in multi-site neuroimaging analyses. Frontiers in Neurology, 13, 923988. 10.3389/fneur.2022.923988

Beer, J. C., Tustison, N. J., Cook, P. A., Davatzikos, C., Sheline, Y. I., Shinohara, R. T., & Linn, K. A. (2020). Longitudinal ComBat: A method for harmonizing longitudinal multi-scanner imaging data. NeuroImage, 220, 117129. 10.1016/j.neuroimage.2020.117129

Ber, R., Hoffman, D., Hoffman, C., Polat, A., Derazne, E., Mayer, A., & Katorza, E. (2017). Volume of Structures in the Fetal Brain Measured with a New Semiautomated Method. American Journal of Neuroradiology, 38(11), 2193–2198. 10.3174/ajnr.A5349

Bethlehem, R. A. I., Seidlitz, J., White, S. R., Vogel, J. W., Anderson, K. M., Adamson, C., Adler, S., Alexopoulos, G. S., Anagnostou, E., Areces-Gonzalez, A., Astle, D. E., Auyeung, B., Ayub, M., Bae, J., Ball, G., Baron-Cohen, S., Beare, R., Bedford, S. A., Benegal, V., … Alexander-Bloch, A. F. (2022). Brain charts for the human lifespan. Nature, 604(7906), 525–533. 10.1038/s41586-022-04554-y

Bozek, J., Griffanti, L., Lau, S., & Jenkinson, M. (2023). Normative models for neuroimaging markers: Impact of model selection, sample size and evaluation criteria. Neuroimage, 268, 119864. 10.1016/j.neuroimage.2023.119864

Chen, A. A., Beer, J. C., Tustison, N. J., Cook, P. A., Shinohara, R. T., Shou, H., & The Alzheimer’s Disease Neuroimaging Initiative. (2022). Mitigating site effects in covariance for machine learning in neuroimaging data. Human Brain Mapping, 43(4), 1179–1195. 10.1002/hbm.25688

Chen, A. A., Luo, C., Chen, Y., Shinohara, R. T., & Shou, H. (2022). Privacy-preserving harmonization via distributed ComBat. NeuroImage, 248, 118822. 10.1016/j.neuroimage.2021.118822

Cole, T. J. (2012). The development of growth references and growth charts. Annals of Human Biology, 39(5), 382–394. 10.3109/03014460.2012.694475

Córdova-Palomera, A., Van Der Meer, D., Kaufmann, T., Bettella, F., Wang, Y., Alnæs, D., Doan, N. T., Agartz, I., Bertolino, A., Buitelaar, J. K., Coynel, D., Djurovic, S., Dørum, E. S., Espeseth, T., Fazio, L., Franke, B., Frei, O., Håberg, A., Le Hellard, S., … Westlye, L. T. (2021). Genetic control of variability in subcortical and intracranial volumes. Molecular Psychiatry, 26(8), 3876–3883. 10.1038/s41380-020-0664-1

De Boer, A. A. A., Bayer, J. M. M., Kia, S. M., Rutherford, S., Zabihi, M., Fraza, C., Barkema, P., Westlye, L. T., Andreassen, O. A., Hinne, M., Beckmann, C. F., & Marquand, A. (2024). Non-Gaussian normative modelling with hierarchical Bayesian regression. Imaging Neuroscience, 2, 1–36. 10.1162/imag_a_00132

Dinsdale, N. K., Jenkinson, M., & Namburete, A. I. L. (2021). Deep learning-based unlearning of dataset bias for MRI harmonisation and confound removal. NeuroImage, 228, 117689. 10.1016/j.neuroimage.2020.117689

Dong, H.-M., Castellanos, F. X., Yang, N., Zhang, Z., Zhou, Q., He, Y., Zhang, L., Xu, T., Holmes, A. J., Thomas Yeo, B. T., Chen, F., Wang, B., Beckmann, C., White, T., Sporns, O., Qiu, J., Feng, T., Chen, A., Liu, X., … Zuo, X.-N. (2020). Charting brain growth in tandem with brain templates at school age. Science Bulletin, 65(22), 1924–1934. 10.1016/j.scib.2020.07.027

Fortin, J.-P., Cullen, N., Sheline, Y. I., Taylor, W. D., Aselcioglu, I., Cook, P. A., Adams, P., Cooper, C., Fava, M., McGrath, P. J., McInnis, M., Phillips, M. L., Trivedi, M. H., Weissman, M. M., & Shinohara, R. T. (2018). Harmonization of cortical thickness measurements across scanners and sites. NeuroImage, 167, 104–120. 10.1016/j.neuroimage.2017.11.024

Fortin, J.-P., Parker, D., Tunç, B., Watanabe, T., Elliott, M. A., Ruparel, K., Roalf, D. R., Satterthwaite, T. D., Gur, R. C., Gur, R. E., Schultz, R. T., Verma, R., & Shinohara, R. T. (2017). Harmonization of multi-site diffusion tensor imaging data. NeuroImage, 161, 149–170. 10.1016/j.neuroimage.2017.08.047

Fortin, J.-P., Sweeney, E. M., Muschelli, J., Crainiceanu, C. M., & Shinohara, R. T. (2016). Removing inter-subject technical variability in magnetic resonance imaging studies. NeuroImage, 132, 198–212. 10.1016/j.neuroimage.2016.02.036

Gage, A. T., Stone, J. R., Wilde, E. A., McCauley, S. R., Welsh, R. C., Mugler, J. P., Tustison, N., Avants, B., Whitlow, C. T., Lancashire, L., Bhatt, S. D., & Haas, M. (2024). Normative Neuroimaging Library: Designing a Comprehensive and Demographically Diverse Dataset of Healthy Controls to Support Traumatic Brain Injury Diagnostic and Therapeutic Development. Journal of Neurotrauma, 41(23–24), 2497–2512. 10.1089/neu.2024.0128

Gardner, M., Shinohara, R. T., Bethlehem, R. A. I., Romero-Garcia, R., Warrier, V., Dorfschmidt, L., Lifespan Brain Chart Consortium, Shanmugan, S., Thompson, P., Seidlitz, J., Alexander-Bloch, A. F., & Chen, A. A. (2024). ComBatLS: A location- and scale-preserving method for multi-site image harmonization. 10.1002/hbm.70197

Hong, J., Hwang, J., & Lee, J.-H. (2023). General psychopathology factor (p-factor) prediction using resting-state functional connectivity and a scanner-generalization neural network. Journal of Psychiatric Research, 158, 114–125. 10.1016/j.jpsychires.2022.12.037

Hu, F., Chen, A. A., Horng, H., Bashyam, V., Davatzikos, C., Alexander-Bloch, A., Li, M., Shou, H., Satterthwaite, T. D., Yu, M., & Shinohara, R. T. (2023). Image harmonization: A review of statistical and deep learning methods for removing batch effects and evaluation metrics for effective harmonization. NeuroImage, 274, 120125. 10.1016/j.neuroimage.2023.120125

Johnson, W. E., Li, C., & Rabinovic, A. (2007). Adjusting batch effects in microarray expression data using empirical Bayes methods. Biostatistics, 8(1), 118–127. 10.1093/biostatistics/kxj037

Klein, A., & Tourville, J. (2012). 101 Labeled Brain Images and a Consistent Human Cortical Labeling Protocol. Frontiers in Neuroscience, 6. 10.3389/fnins.2012.00171

Koenker, R., & Bassett, G. (1978). Regression Quantiles. Econometrica, 46(1), 33–50. 10.2307/1913643

Kruggel, F., Turner, J., & Muftuler, L. T. (2010). Impact of scanner hardware and imaging protocol on image quality and compartment volume precision in the ADNI cohort. NeuroImage, 49(3), 2123–2133. 10.1016/j.neuroimage.2009.11.006

Moyer, D., Ver Steeg, G., Tax, C. M. W., & Thompson, P. M. (2020). Scanner invariant representations for diffusion MRI harmonization. Magnetic Resonance in Medicine, 84(4), 2174–2189. 10.1002/mrm.28243

Nobis, L., Manohar, S. G., Smith, S. M., Alfaro-Almagro, F., Jenkinson, M., Mackay, C. E., & Husain, M. (2019). Hippocampal volume across age: Nomograms derived from over 19,700 people in UK Biobank. NeuroImage: Clinical, 23, 101904. 10.1016/j.nicl.2019.101904

Pomponio, R., Erus, G., Habes, M., Doshi, J., Srinivasan, D., Mamourian, E., Bashyam, V., Nasrallah, I. M., Satterthwaite, T. D., Fan, Y., Launer, L. J., Masters, C. L., Maruff, P., Zhuo, C., Völzke, H., Johnson, S. C., Fripp, J., Koutsouleris, N., Wolf, D. H., … Davatzikos, C. (2020). Harmonization of large MRI datasets for the analysis of brain imaging patterns throughout the lifespan. NeuroImage, 208, 116450. 10.1016/j.neuroimage.2019.116450

Reig, S., Sánchez-González, J., Arango, C., Castro, J., González-Pinto, A., Ortuño, F., Crespo-Facorro, B., Bargalló, N., & Desco, M. (2009). Assessment of the increase in variability when combining volumetric data from different scanners. Human Brain Mapping, 30(2), 355–368. 10.1002/hbm.20511

Rutherford, S., Barkema, P., Tso, I. F., Sripada, C., Beckmann, C. F., Ruhe, H. G., & Marquand, A. F. (2023). Evidence for embracing normative modeling. eLife, 12, e85082. 10.7554/eLife.85082

Solanes, A., Palau, P., Fortea, L., Salvador, R., González-Navarro, L., Llach, C. D., Valentí, M., Vieta, E., & Radua, J. (2021). Biased accuracy in multisite machine-learning studies due to incomplete removal of the effects of the site. Psychiatry Research: Neuroimaging, 314, 111313. 10.1016/j.pscychresns.2021.111313

Thompson, P. M., Jahanshad, N., Ching, C. R. K., Salminen, L. E., Thomopoulos, S. I., Bright, J., Baune, B. T., Bertolín, S., Bralten, J., Bruin, W. B., Bülow, R., Chen, J., Chye, Y., Dannlowski, U., De Kovel, C. G. F., Donohoe, G., Eyler, L. T., Faraone, S. V., Favre, P., … for the ENIGMA Consortium. (2020). ENIGMA and global neuroscience: A decade of large-scale studies of the brain in health and disease across more than 40 countries. Translational Psychiatry, 10(1), 100. 10.1038/s41398-020-0705-1

Tustison, N. J., Holbrook, A. J., Avants, B. B., Roberts, J. M., Cook, P. A., Reagh, Z. M., Duda, J. T., Stone, J. R., Gillen, D. L., Yassa, M. A., & Alzheimer’s Disease Neuroimaging Initiative. (2019). Longitudinal Mapping of Cortical Thickness Measurements: An Alzheimer’s Disease Neuroimaging Initiative-Based Evaluation Study. Journal of Alzheimer’s Disease: JAD, 71(1), 165–183. 10.3233/JAD-190283

Yamashita, A., Yahata, N., Itahashi, T., Lisi, G., Yamada, T., Ichikawa, N., Takamura, M., Yoshihara, Y., Kunimatsu, A., Okada, N., Yamagata, H., Matsuo, K., Hashimoto, R., Okada, G., Sakai, Y., Morimoto, J., Narumoto, J., Shimada, Y., Kasai, K., … Imamizu, H. (2019). Harmonization of resting-state functional MRI data across multiple imaging sites via the separation of site differences into sampling bias and measurement bias. PLOS Biology, 17(4), e3000042. 10.1371/journal.pbio.3000042

Yu, M., Linn, K. A., Cook, P. A., Phillips, M. L., McInnis, M., Fava, M., Trivedi, M. H., Weissman, M. M., Shinohara, R. T., & Sheline, Y. I. (2018). Statistical harmonization corrects site effects in functional connectivity measurements from multi-site fMRI data. Human Brain Mapping, 39(11), 4213–4227. 10.1002/hbm.24241

Zhang, R., Oliver, L. D., Voineskos, A. N., & Park, J. Y. (2023). RELIEF: A structured multivariate approach for removal of latent inter-scanner effects. Imaging Neuroscience, 1, 1–16. 10.1162/imag_a_00011

Zhu, A. H., Nir, T. M., Javid, S., Villalón-Reina, J. E., Rodrigue, A. L., Strike, L. T., De Zubicaray, G. I., McMahon, K. L., Wright, M. J., Medland, S. E., Blangero, J., Glahn, D. C., Kochunov, P., Williamson, D. E., Håberg, A. K., Thompson, P. M., & Jahanshad, N. (2025). Lifespan reference curves for harmonizing multi-site regional brain white matter metrics from diffusion MRI. Scientific Data, 12(1), 748. 10.1038/s41597-025-05028-2

Zhu, J.-Y., Park, T., Isola, P., & Efros, A. A. (2017). Unpaired Image-to-Image Translation Using Cycle-Consistent Adversarial Networks. 2017 IEEE International Conference on Computer Vision (ICCV), 2242–2251. 10.1109/ICCV.2017.244

